# Optineurin promotes aggregation of mutant huntingtin and mutant ataxin-3, and reduces cytotoxicity of aggregates

**DOI:** 10.1101/2020.08.13.249201

**Authors:** Shivranjani C Moharir, Akhouri Kishore Raghawan, Ghanshyam Swarup

**Affiliations:** CSIR-Centre for Cellular and Molecular Biology, Hyderabad-500007, India

**Author notes:** **Correspondence to:** Ghanshyam Swarup, CSIR-Centre for Cellular and Molecular Biology, Hyderabad-500007, India.

**Keywords:** Optineurin (OPTN), mutant protein aggregates, neurodegeneration, huntingtin, ataxin-3

## Abstract

Optineurin (OPTN), a cytoplasmic adaptor protein involved in cargo selective autophagy of bacteria, damaged mitochondria and mutant protein aggregates, is frequently seen in pathological structures containing protein aggregates, associated with several neurodegenerative diseases. However, the function of OPTN in these protein aggregates is not known. Here, we have explored the role of OPTN in mutant protein aggregation and in cytoprotection from toxicity of mutant proteins. Mutant huntingtin (mHtt) and mutant ataxin-3 (mAtax-3) showed reduced formation of aggregates in *Optn*^*−/−*^ mouse embryonic fibroblasts as compared with wild type cells. Co-expression of OPTN enhanced aggregate formation by mHtt and mAtax-3 in *Optn*^*−/−*^ cells. C-terminal domain of OPTN (412-577 amino acids) was necessary and sufficient to promote aggregate formation by these mutant proteins. The E478G mutant of OPTN, defective in ubiquitin-binding and autophagy, was also able to promote aggregation of mHtt and mAtax-3. OPTN and its C-terminal domain form a complex with the chaperone HSP70 known to promote mutant protein aggregation. Overexpression of mHtt or mAtax-3 induced more cell death in *Optn*^*−/−*^ cells compared with wild type cells. Importantly, compared to wild type cells, Optn-deficient cells having mHtt or mAtax-3 aggregates showed higher level of cell death in neuronal (N2A) and non-neuronal cells. Our results show that OPTN promotes formation of mutant huntingtin and mutant ataxin-3 aggregates, and this function of OPTN might be mediated through interaction with HSP70 chaperones. Our results also show that OPTN reduces cytotoxicity caused by these mutant protein aggregates.

**Significance statement:** The hallmark of several neurodegenerative diseases like amyotrophic lateral sclerosis, Huntington’s disease, Parkinson’s disease, Alzheimer’s disease and Pick’s disease is the formation of pathological structures containing aggregated proteins, and OPTN is frequently observed in these structures. What role optineurin plays in those aggregates is not clear. Our results show that OPTN promotes aggregation of mutant huntingtin and mutant ataxin-3, and reduces cytotoxicity of aggregates in neuronal and non-neuronal cells. We suggest that OPTN provides cytoprotection in three different ways-by promoting mutant protein aggregation, by reducing cytotoxicity of aggregates and by autophagy-dependent clearance of aggregates reported earlier. These properties of OPTN provide a possible explanation for its association with various pathological structures containing protein aggregates seen in several neurodegenerative diseases.

## Introduction

Formation of cytoplasmic or nuclear protein aggregates in neurons is the hallmark of several neurodegenerative diseases. The aggregating proteins either have point mutations in their sequences as in case of *SOD1* (1, 2), *FUS* (3, 4), *TDP43* (5, 6) and *OPTN* (7) mutants causing amyotrophic lateral sclerosis (ALS), or they have abnormally long poly-glutamine sequences originating from CAG repeats in the DNA, as in case of huntingtin protein causing Huntington’s disease, and ataxin-3 causing spinocerebellar ataxia type-3 (8, 9). Mutated misfolded proteins are not just non-functional, if not cleared from cells they can interfere with normal cellular processes (8). These mutant proteins are either present in non-aggregated form in the cytoplasm or form compartmentalized inclusion bodies or aggregates in the cytoplasm or nucleus. Protein aggregates are usually seen in the late stages of diseases (10). Several reports suggest that aggregates are associated with toxicity to the cells (11, 12). However, the exact nature and physical properties of the aggregates or their precursors, which are responsible for causing toxicity, is still under dispute. Emerging evidences suggest that the monomers and the intermediate fibrillary structures of mutant proteins non-aggregated in cytoplasm are more toxic to cells than the aggregates or inclusion bodies (13–16). Some studies actually show that inclusion body formation improves neuronal survival and decreases the levels of mutant protein elsewhere in the neurons (17–19). The conditions that suppress formation of inclusion bodies of mutant proteins result in higher cell death (14, 20, 21).

The mutant protein aggregates are ubiquitin positive and are cleared by the process of autophagy (22, 23), a catabolic ‘waste disposal process’ deployed by the cells to clear mutant aggregated proteins and damaged organelles. The cargo to be degraded is recognized with the help of designated ‘autophagy receptors’, which recruit the cargo to the autophagosomes. The autophagosomes fuse with lysosomes to form autolysosomes where the actual degradation process takes place (24, 25). Perturbation of autophagy is associated with neurodegeneration and other disorders (26). Optineurin (OPTN) is a cytoplasmic adaptor protein involved in cargo selective autophagy of certain bacteria, damaged mitochondria and mutant protein aggregates that are associated with neurodegenerative diseases (27–30). Mutations in OPTN are associated with neurodegenerative diseases such as glaucoma and ALS (7, 31–33). An important feature of several neurodegenerative diseases like ALS, Huntington’s disease, Parkinson’s disease, Alzheimer’s disease, Pick’s disease and Creutzfeld-Jacob disease is the formation of pathological structures containing aggregated proteins, and OPTN is frequently observed in these structures (7, 34, 35). What role optineurin plays in those aggregates is yet to be explored. The autophagy receptor function of OPTN is involved in clearance of mutant huntingtin (with expanded poly-glutamine repeats) and mutant SOD1 protein aggregates (29, 30, 36). A study has shown that medium projection neurons, a subset of striatal neurons, which express lower level of optineurin, are more vulnerable in Huntington’s disease mouse model as compared to interneurons which express optineurin abundantly (37). Thus, presence of OPTN might have some protective role in neurodegenerative diseases caused by mutant proteins.

Chaperones of HSP70 family have been implicated in influencing aggregation of mutant proteins (38, 39). HSC70 and its co-chaperones have been shown to suppress mutant protein aggregation (38, 40) but the role of HSP70 in mutant protein aggregation is controversial. Some studies have shown that HSP70 suppresses mutant protein aggregation whereas other reports show that HSP70 promotes aggregation. Genetic studies in yeast have shown that cytosolic HSP70 (Ssa family) is required for mutant Huntingtin aggregation (41). Recent studies suggest that aggregation of mutant proteins to form inclusion bodies is facilitated by chaperones of HSP70 family that leads to reduction of diffusible mutant protein (which is more toxic) resulting in enhanced cell survival (42).

In this study, we have explored the role of optineurin in mutant protein aggregation and in cytoprotection from toxicity of mutant proteins. We have used two mutant protein models, a short fragment of huntingtin protein having 97 glutamine repeats (mHtt) (36) and, a short fragment of ataxin-3 protein having 130 glutamine repeats (mAtax-3) (43) to understand the role of optineurin in protein aggregation and cytoprotection. By using *Optn*^*−/−*^ and wild type (WT) mouse embryonic fibroblasts (MEFs), we show that optineurin promotes formation of mHtt 97Q and mAtax-3 130Q aggregates. Further, we found that C-terminal sequence of OPTN consisting of 412-577 amino acids was necessary as well as sufficient for promoting aggregation of these mutant proteins. Optineurin interacts with HSP70 and C-terminal domain of optineurin is sufficient for this interaction. We also observed that optineurin protects neuronal as well as non-neuronal cells from cytotoxicity caused by aggregates.

## Results

### Optineurin promotes aggregation of mutant huntingtin and mutant ataxin-3

Optineurin mediates autophagy dependent clearance of protein aggregates formed by mHtt and some other mutant proteins associated with neurodegenerative diseases in neuronal and non-neuronal cells (29, 30). Recently we have generated *Optn*^*−/−*^ mouse embryonic fibroblasts (MEFs) which show reduced non-selective (basal and starvation induced) autophagy as seen by reduced formation of LC3-II, and lower number of autophagosomes and autolysosomes in comparison with wild type cells (36, 44). We examined the formation of mHtt aggregates in *Optn*^*−/−*^ and wild type MEFs by expressing GFP tagged mHtt and observing under a fluorescence microscope. We analysed the percentage of GFP expressing cells forming aggregates. Contrary to our expectation, we observed that compared to wild type cells, *Optn*^*−/−*^ MEFs showed significantly lower percentage of cells forming mHtt aggregates (Fig. 1 A and B). The expression level of GFP-mHtt 97Q was comparable in optineurin-deficient cells and wild type cells as seen by western blot (Fig. 1C). We also examined the aggregation of mHtt by determining its level in detergent insoluble and soluble fractions by western blotting. Quantification of western blots revealed that the detergent insoluble fraction of mHtt was significantly higher in WT cells in comparison with Optn-deficient cells (Fig. 1D and E). These results suggested that optineurin promotes aggregation of mHtt in MEFs. We examined the role of optineurin in aggregate formation using another protein, mutant ataxin-3, which also has expanded poly glutamine repeats. Expression of GFP tagged mAtax-3 in *Optn*^*−/−*^ cells resulted in reduced formation of aggregates in comparison with wild type cells (Fig. 1F and G). The expression of GFP-mAtax-3 was similar in wild type and optineurin deficient cells (Fig. 1H). The detergent insoluble fraction of mHtt was significantly higher in WT cells in comparison with Optn-deficient cells (Fig. 1I and J). Co-expression of HA-tagged wild type OPTN resulted in enhanced formation of mHtt and mAtax-3 aggregates in *Optn*^*−/−*^ cells (Fig. 2C-G). These results suggest that endogenous as well as overexpressed optineurin promotes aggregation of mHtt and mAtax-3. These results also indicate that optineurin mediated autophagy is not able to clear mHtt and mAtax-3 aggregates in MEFs.

**Figure 1.**
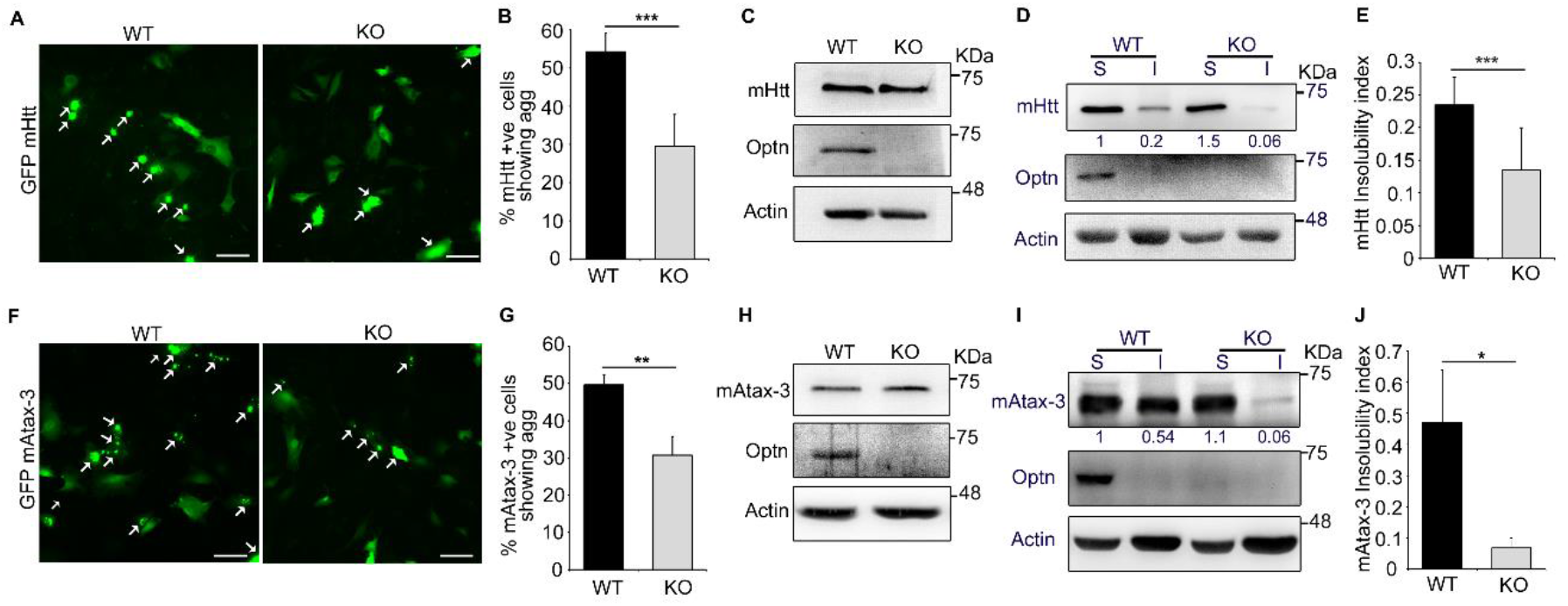
Optineurin promotes aggregation of mHtt: **(A)** Microscopy images showing GFP-mHtt expressing cells forming aggregates in WT and *Optn*^*−/−*^ (KO) MEFs after 48 hours of transfection. Arrows indicate cells showing aggregates of mHtt. Scale bar: 100 μm. **(B)** Bar diagram showing the percentage of GFP-mHtt expressing cells with aggregates (agg) in WT and *Optn*^*−/−*^ MEFs. n= 5. Bars represent mean ± SD. ***p ≤ 0.001. **(C)** Western blot showing expression of GFP-mHtt in WT and *Optn*^*−/−*^ MEFs. **(D)** Western blot analysis of NP-40 lysis buffer soluble and insoluble fractions of WT and *Optn*^*−/−*^ MEFs transfected with GFP-mHtt for 48 hours. **(E)** Bar diagram showing insolubility index of GFP-mHtt in WT and *Optn*^*−/−*^ MEFs. n=6. ***p ≤ 0.001. **(F)** Microscopy images showing GFP-mAtax-3 expressing cells forming aggregates in WT and *Optn*^*−/−*^ MEFs after 48 hours of transfection. Arrows indicate cells showing aggregates of mAtax-3. Scale bar: 100 μm. **(G)** Bar diagram showing the percentage of GFP-mAtax-3 expressing cells with aggregates in WT and *Optn*^*−/−*^ MEFs. n= 3. **p ≤ 0.01. **(H)** Western blot showing expression of GFP-mAtax-3 in WT and *Optn*^*−/−*^ MEFs. **(I)** Western blot analysis of NP-40 lysis buffer soluble and insoluble fractions of WT and *Optn*^*−/−*^ MEFs transfected with GFP-mAtax-3 for 48 hours. **(J)** Bar diagram showing insolubility index of GFP-mAtax-3 in WT and *Optn*^*−/−*^ MEFs. n=3. *p ≤ 0.05.

**Figure 2:**
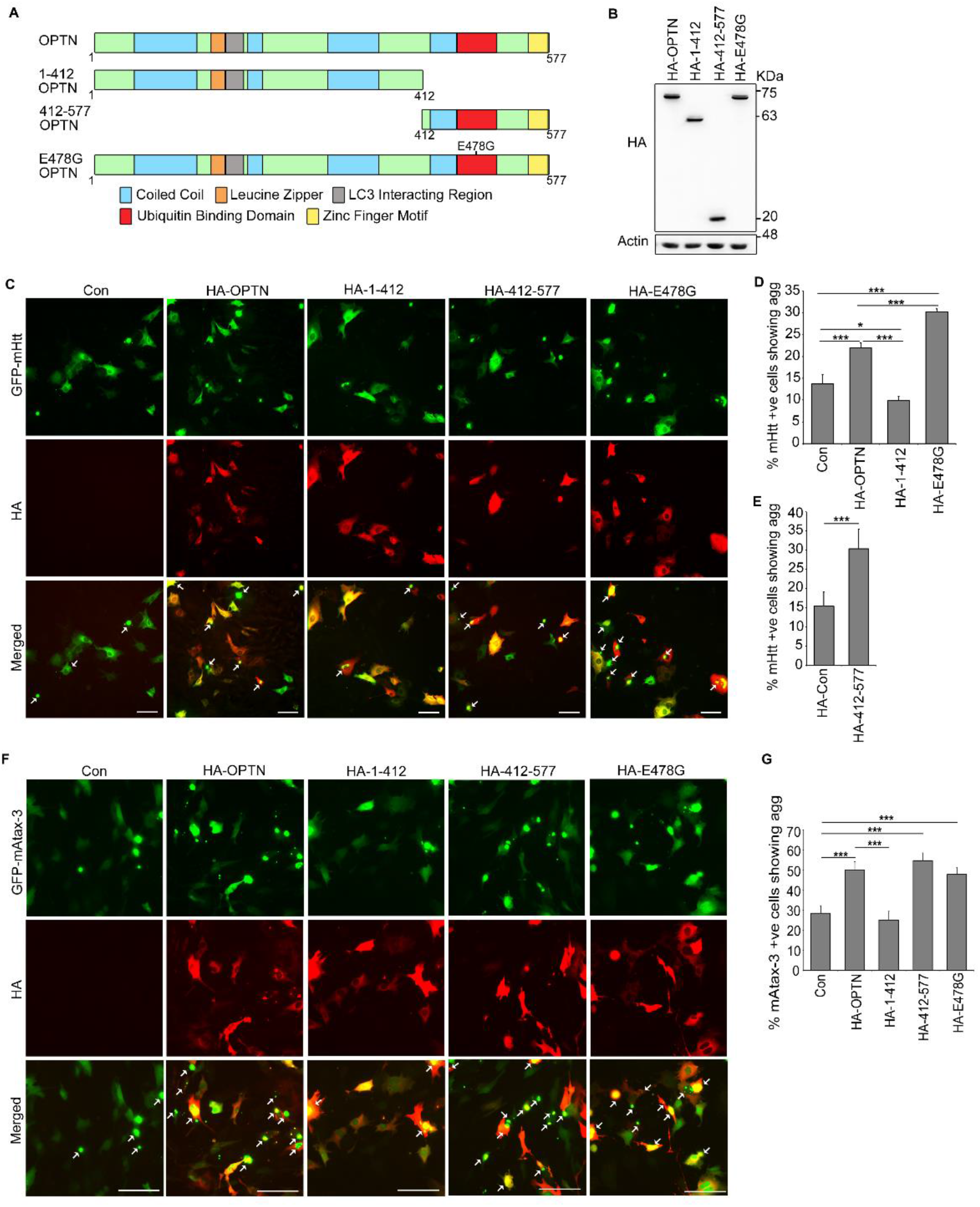
C-terminal region of OPTN is sufficient to promote aggregation of mHtt and mAtax-3: **(A)** Schematic representation of domain structure of OPTN, 1-412-OPTN, 412-577-OPTN and E478G-OPTN. **(B)** Western blot showing expression of HA-OPTN, HA-1-412-OPTN, HA-412-577-OPTN and HA-E478G-OPTN in *Optn*^*−/−*^ MEFs. The blot was probed with HA antibody. **(C and F)** Microscopy images showing the effect of expression of HA-OPTN and its mutants on aggregation of GFP-mHtt **(C)** and mAtax-3 **(F)** in *Optn*^*−/−*^ MEFs. Arrows indicate cells showing aggregates of mHtt. Scale bar: 100μm. **(D)** and **(E)** Bar diagrams showing the effect of expression of HA-OPTN and its mutants on percentage of GFP-mHtt and positive cells forming aggregates (agg) in *Optn*^*−/−*^ MEFs. n= 6 for HA-412-577-OPTN. For all others, n=4. Bars represent mean ± SD. ***p ≤ 0.001 and *p ≤ 0.05. **(G)** Bar diagram showing the effect of expression of HA-OPTN and its mutants on percentage of GFP-mAtax-3 positive cells forming aggregates in *Optn*^*−/−*^ MEFs. n= 6. ***p ≤ 0.001.

### C-terminal domain of optineurin is sufficient to promote mutant protein aggregation

Optineurin is an adaptor protein, which has coiled coil domains, an LIR (LC3-interacting region), a UBD (ubiquitin binding domain) and a zinc finger domain (Fig. 2A) (32). We determined the domain requirement of OPTN for promoting aggregation of mutant proteins by using various deletion and point mutants (Fig. 2 A and B). GFP-mHtt was co-expressed with HA tagged OPTN or its mutants in *Optn*^*−/−*^ cells and aggregate formation was analysed in cells expressing both proteins. OPTN construct (1-412aa) lacking C-terminal domain was unable to enhance the percentage of cells forming mHtt aggregates in *Optn*^*−/−*^ cells (Fig. 2C and D). Surprisingly, C-terminal domain (412-577aa) of OPTN was able to enhance the percentage of cells forming mHtt aggregates as efficiently as wild type OPTN (Fig. 2C and E). Since C-terminal domain of OPTN includes the UBD and mHtt aggregates are known to be ubiquitinated, we examined the requirement of UBD by using a mutant of OPTN, E478G that is defective in ubiquitin-binding and autophagy. Co-expression of E478G mutant resulted in enhanced formation of mHtt aggregates in *Optn*^*−/−*^ cells (Fig. 2C and D). In fact, compared to wild type OPTN, the E478G mutant, which is observed in inclusion bodies in ALS patient samples (7), was more effective in enhancing aggregate formation by mHtt (Fig. 2C and D). We also examined the domain requirement of OPTN for enhancing mAtax-3 aggregation and observed that C-terminal domain of OPTN was necessary and sufficient to promote mAtax-3 aggregate formation (Fig. 2F and G). E478G mutant of OPTN was as effective in promoting mAtax-3 aggregate formation as wild type OPTN (Fig. 2F and G). The expression of various mutants of OPTN was similar as determined by western blot (Fig. 2B). These results suggest that binding of OPTN to ubiquitin is not required to promote aggregation of mutant proteins. These results also indicate that promoting the formation of mutant protein aggregates and their clearance by autophagy are two separate functions of OPTN.

### OPTN localizes with the aggregates and forms a shell around the aggregates through C-terminal domain

Since deficiency of endogenous optineurin in *Optn*^*−/−*^ MEFs resulted in reduced formation of mHtt and mAtax-3 aggregates, we examined whether endogenous optineurin goes to the mutant protein aggregates. Using fluorescent confocal microscope, we observed that, endogenous optineurin was localized to the periphery and formed a shell around large aggregates of mHtt as well as mAtax-3, while in case of smaller aggregates, optineurin localized with mHtt and mAtax-3 throughout the aggregate (Fig. 3A and B). Overexpressed OPTN also formed a shell around mHtt and mAtax-3 aggregates and showed localization at the periphery of these aggregates (Fig. 3C and Supplementary figure S1A). The OPTN mutants E478G and 412-577-OPTN, which aided in aggregation of mutant proteins, also localized with mHtt and mAtax-3 aggregates at the periphery and formed the characteristic shell around them (Fig. 3C and Supplementary figure S1A). The mutant 1-412-OPTN, which did not promote aggregation of mHtt and mAtax-3, did not localize with the aggregates and it did not form a shell around the aggregates (Fig. 3C and Supplementary figure S1A). These results show that C-terminal domain of OPTN is necessary and sufficient for localization around mutant protein aggregates. To rule out the possibility that formation of shell around the aggregates is an artifact of immunostaining, we co-expressed mCherry-mHtt and GFP-Optn and observed that like endogenous optineurin and HA-OPTN, GFP-Optn also formed a shell around mCherry-mHtt (Supplementary figure S1B).

**Figure 3:**
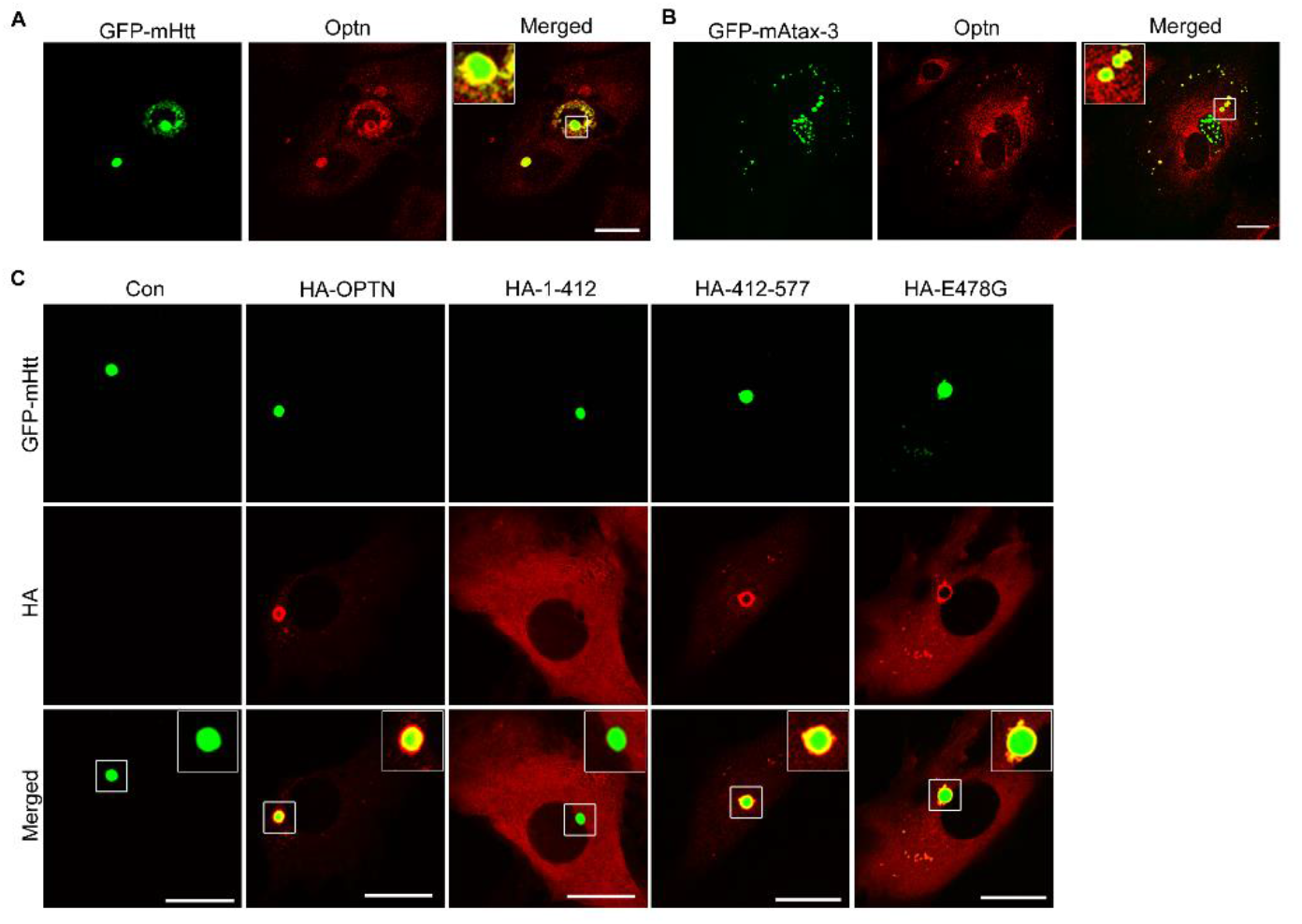
C-terminal domain of optineurin is necessary and sufficient to form a shell around mutant protein aggregates: Confocal microscopy images showing endogenous optineurin forming a shell around GFP-mHtt **(A)** and GFP-mAtax-3 **(B)** aggregates in WT MEFs. Three confocal sections are merged in the images. Scale bar: 25μm. **(C)** Confocal microscopy images showing that HA-OPTN, HA-412-577-OPTN and E478G-OPTN form a shell around GFP-mHtt aggregates in *Optn*^*−/−*^ MEFs whereas HA-1-412-OPTN does not. Scale bar: 25μm. Three confocal sections are merged in the images.

### Optineurin promotes aggregation of endogenous proteins

To understand the role of optineurin in modulating aggregation of endogenous misfolded proteins, we used p62 as the marker for endogenous protein aggregates (45) and counted the percentage of cells showing p62-positive puncta in WT and *Optn*^*−/−*^ MEFs. WT MEFs showed higher percentage of cells showing p62-positive puncta as compared to KO MEFs (Fig. 4A and B). The level of p62 protein was similar in both WT and *Optn*^*−/−*^ MEFs (Fig. 4C and D). This suggested that in *Optn*^*−/−*^ MEFs, the basal level of formation of p62-positive puncta is compromised. Next, we blocked autophagic degradation of proteins by using chloroquine in WT and *Optn*^*−/−*^ MEFs and counted the percentage of cells showing p62-positive puncta. We observed that upon treatment with chloroquine, *Optn*^*−/−*^ MEFs showed significantly lower percentage of cells showing p62-positive puncta (Fig. 4A and B). This indicated that optineurin promotes aggregation of endogenous misfolded proteins which are degraded by autophagy. Over expression of OPTN in *Optn*^*−/−*^ MEFs increased the percentage of cells showing p62 puncta as compared to control group (Fig. 4E and F). Many of the over-expressed OPTN puncta were positive for p62 and most of the over-expressed OPTN puncta were positive for ubiquitin (Fig 4G and H). In WT MEFs, p62 puncta co-localized with ubiquitin puncta (Fig. 4I).

**Figure 4:**
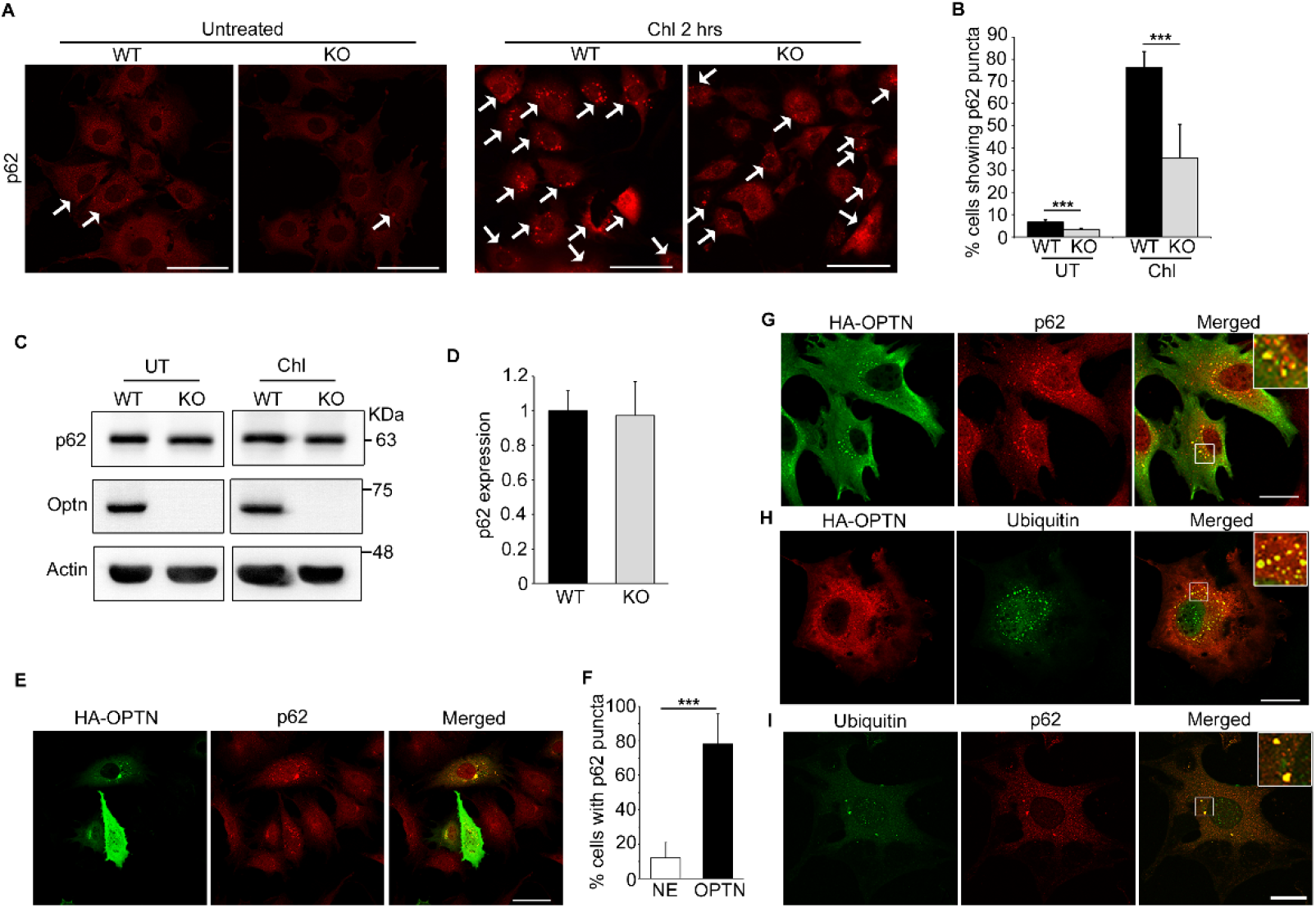
Optineurin promotes aggregation of endogenous proteins: **(A)** Microscopy images showing percentage of cells having p62 puncta in WT and *Optn*^*−/−*^ MEFs in cells left untreated (UT) or treated with 50 μM chloroquine for 2 hrs. Arrows indicate cells showing p62 puncta. Scale bar: 100μm. **(B)** Bar diagram showing percentage of cells having p62 puncta in WT and *Optn*^*−/−*^ MEFs at basal level or upon treatment with 50 μM chloroquine for 2 hrs. n=6. Bars represent mean ± SD. ***p ≤ 0.001. **(C)** Western blot showing p62 levels in WT and *Optn*^*−/−*^ MEFs. **(D)** Bar diagram showing p62 levels in WT and *Optn*^*−/−*^ MEFs. n=5. **(E)** Microscopy images showing that expression of HA-OPTN in *Optn*^*−/−*^ MEFs increases p62 puncta. Scale bar: 50μm. **(F)** Bar diagram showing percentage of *Optn*^*−/−*^ MEFs having p62 puncta upon expression of HA-OPTN. n=3. **(G)** Confocal microscopy images showing co-localization of HA-OPTN and endogenous p62 in *Optn*^*−/−*^ MEF. Scale bar: 25μm. **(H)** Confocal microscopy images showing co-localization of HA-OPTN and endogenous ubiquitin in *Optn*^*−/−*^ MEF. Scale bar: 25μm. **(I)** Confocal microscopy images showing co-localization of endogenous ubiquitin and p62 in WT MEF. Scale bar: 25μm. Three confocal sections were merged in the images.

### How does optineurin promote mutant protein aggregation?

We attempted to understand the molecular mechanism of optineurin promoted mutant protein aggregation. Molecular chaperones of HSP family (HSC70, HSP70, HSP90) are known to affect mutant protein aggregation. HSC70 has been shown to suppress protein aggregation whereas HSP70 and heat shock are reported to enhance protein aggregation (38–40, 42). We hypothesized that OPTN may be promoting aggregation of mutant proteins by modulating the function of HSP70 family chaperones. Therefore, we examined the interaction of OPTN with HSP70, HSC70 and HSP90 by immunoprecipitation, and observed that these proteins form a complex with over-expressed OPTN (Fig. 5A). Endogenous OPTN also formed a complex with HSP70 and HSP90 (Fig. 5B). We also analysed the interaction of deletion constructs of OPTN with these proteins in HEK293T cells. HSP90 showed interaction with 412-577-OPTN, which promotes protein aggregation, but not with 1-412-OPTN (Fig. 5C). HSP70 and HSC70 showed enhanced interaction with 412-577-OPTN in comparison with 1-412-OPTN. The level of HSP90 and HSC70 was comparable in WT and Optn-deficient cells, whereas HSP70 level was lower in Optn-deficient cells (Fig. 5D and E). We over-expressed HA-OPTN, HA-1-412-OPTN, HA-412-577-OPTN and E478G-OPTN in *Optn*^*−/−*^ MEFs and observed that none of these mutants affected the levels of HSP70 or HSP90 (Supplementary Figure S2). These results provide some evidence for involvement of HSP70/HSP90 in mediating OPTN promoted protein aggregation. Treatment of cells with 17-AAG (17-N-allylamino-17-demethoxygeldanamycin), a chemical inhibitor of HSP90, resulted in reduced level of mHtt but percentage of cells showing aggregates was increased in both WT and Optn-deficient cells (Fig. 5F, G and H). Treatment of cells with 17-AAG also resulted in induction of HSP70 in both WT and Optn-deficient cells (Fig. 5F). These results indicate that HSP90 is not likely to be involved in mediating optineurin-promoted mHtt aggregation but HSP70 might be involved in this process.

**Figure 5:**
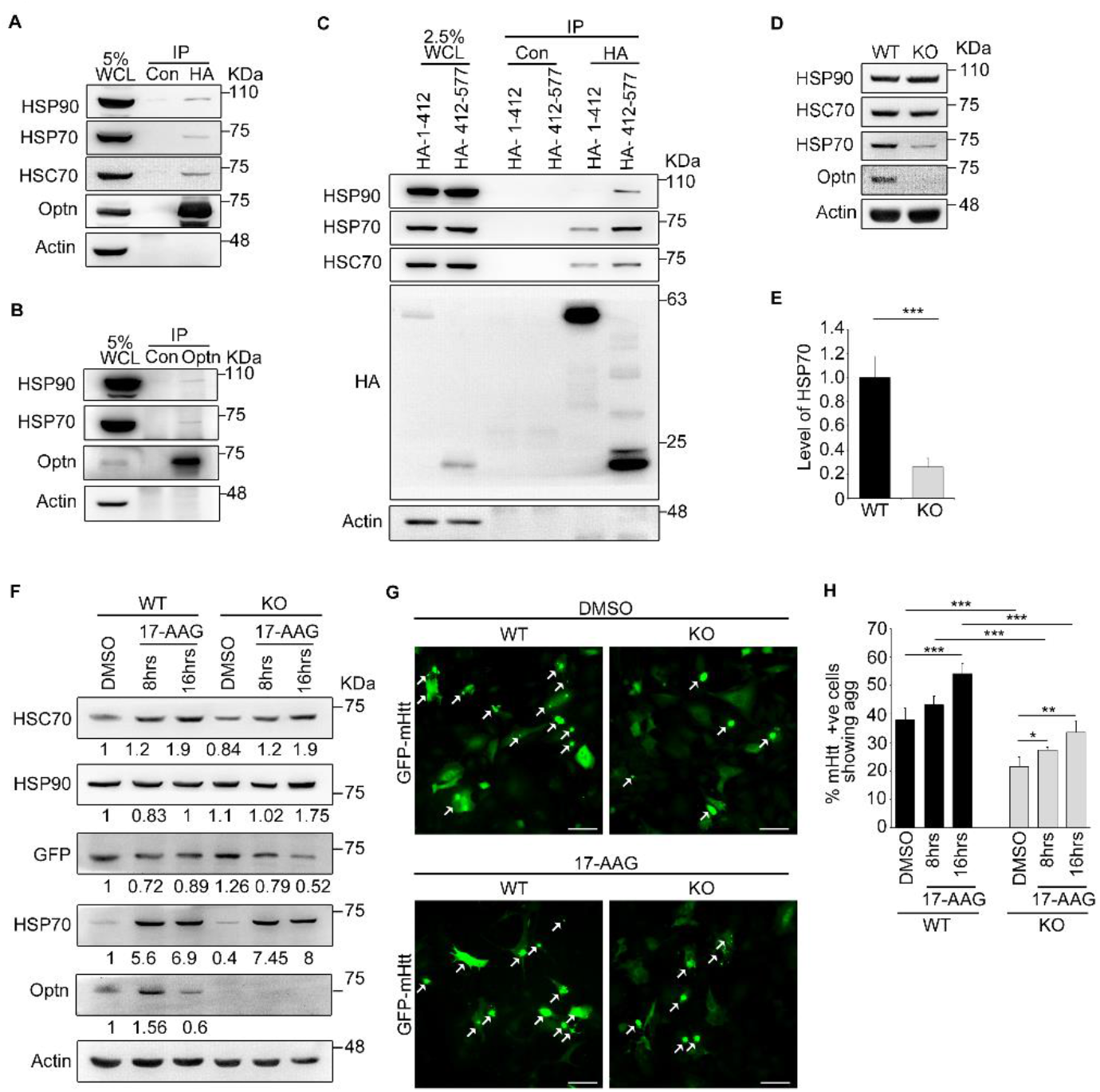
Optineurin interacts with HSP90, HSP70 and HSC70: **(A)** Lysates of HEK293T cells expressing HA-OPTN were subjected to immunoprecipitation using HA antibody agarose beads or normal immunoglobulin (Con) bound to agarose beads as a control, and analyzed for interaction with HSP90, HSP70 and HSC70 by western blotting. HSP70 blot was de-probed and re-probed for HSC70. WCL, whole cell lysates. **(B)** Lysates of HEK293T cells were subjected to immunoprecipitation using optineurin antibody or normal immunoglobulin as a control and analysed for interaction with HSP90 and HSP70 by western blotting. **(C)** Lysates of HEK293T cells expressing HA-1-412-OPTN and HA-412-577-OPTN were subjected to immunoprecipitation using HA antibody agarose beads or normal immunoglobulin bound to agarose beads, and analyzed for interaction with HSP90, HSP70 and HSC70 by western blotting. **(D)** Western blot showing the expression of HSP90, HSP70 and HSC70 in WT and *Optn*^*−/−*^ MEFs. **(E)** Bar diagram showing the level of HSP70 in WT and *Optn*^*−/−*^ MEFs. n=11. Bars represent mean ± SD. ***p ≤ 0.001. **(F)** WT and *Optn*^*−/−*^ MEFs were transfected with GFP-mHtt and left untreated or treated with 0.5μM 17-AAG for the last 8 or 16 hours. Cell lysates were prepared and analysed by western blotting after 48 hours of transfection. **(G)** Microscopy images showing GFP-mHtt expressing cells forming aggregates in WT and *Optn*^*−/−*^ MEFs left untreated or treated with 0.5μM 17-AAG for the last 8 or 16 hours upon transfection with GFP-mHtt for total 48 hours. Arrows indicate cells showing aggregates of mHtt. Scale bar: 100 μm. **(H)** Bar diagram showing the percentage of GFP-mHtt expressing cells with aggregates (agg) in WT and *Optn*^*−/−*^ MEFs left untreated or treated with 0.5μM 17-AAG for 8 or 16 hours. n= 4. *p ≤ 0.05, **p ≤ 0.01 and ***p ≤ 0.001.

### Optineurin reduces the cytotoxicity of mutant protein aggregates

Mutant huntingtin has cytotoxic effect which results in degeneration of neurons that leads to pathogenesis. We examined the effect of optineurin deficiency on cell death induction by mHtt in MEFs. GFP-mHtt was expressed in wild type and *Optn*^*−/−*^ cells and after 48 hours, cell death was determined by staining of cells for externalized phosphatidylserine with Annexin V and nuclear DNA with 7-AAD. Cell death was quantified by microscopic examination. Expression of mHtt resulted in significantly higher level of cell death in *Optn*^*−/−*^ cells compared to wild type cells, suggesting a cytoprotective role for optineurin (Fig. 6A and C). Induction of cell death upon expression of mHtt could be due to cytotoxicity of the aggregates or due to the non-aggregated protein in the cytoplasm. Therefore, we analysed death in cells showing mHtt aggregates and compared it with the death in cells showing non-aggregated mHtt. In wild type cells, aggregated mHtt induced significantly less cell death in comparison with non-aggregated mHtt (Fig 6A and D). However, in optineurin-deficient cells, compared to non-aggregated mHtt, aggregated mHtt induced significantly more cell death (Fig 6D). A comparison of wild type and *Optn*^*−/−*^ cells showed that aggregated mHtt induced significantly more cell death in *Optn*^*−/−*^ cells, whereas cell death induced by non-aggregated mHtt was not affected significantly by deficiency of optineurin (Fig. 6D). These results indicate that optineurin provides protection from cytotoxic effects of mHtt aggregates. Next, we examined the toxicity caused by mAtax-3 protein in WT and *Optn*^*−/−*^ cells. We observed that compared to WT cells, *Optn*^*−/−*^ MEFs were more susceptible to cell death induced by mAtax-3 expression as seen by higher Annexin V and 7-AAD staining (Fig. 6B and E). In both WT and *Optn*^*−/−*^ MEFs, mAtax-3 aggregates were more toxic as compared to non-aggregated mAtax-3 (Fig. 6B and F). A comparison of wild type and *Optn*^*−/−*^ cells showed that aggregated mAtx-3 induced significantly more cell death in *Optn*^*−/−*^ cells (Fig. 6B and F). These results suggest that optineurin deficiency increases cytotoxicity caused by aggregates of mAtax-3 as well as mHtt.

**Figure 6:**
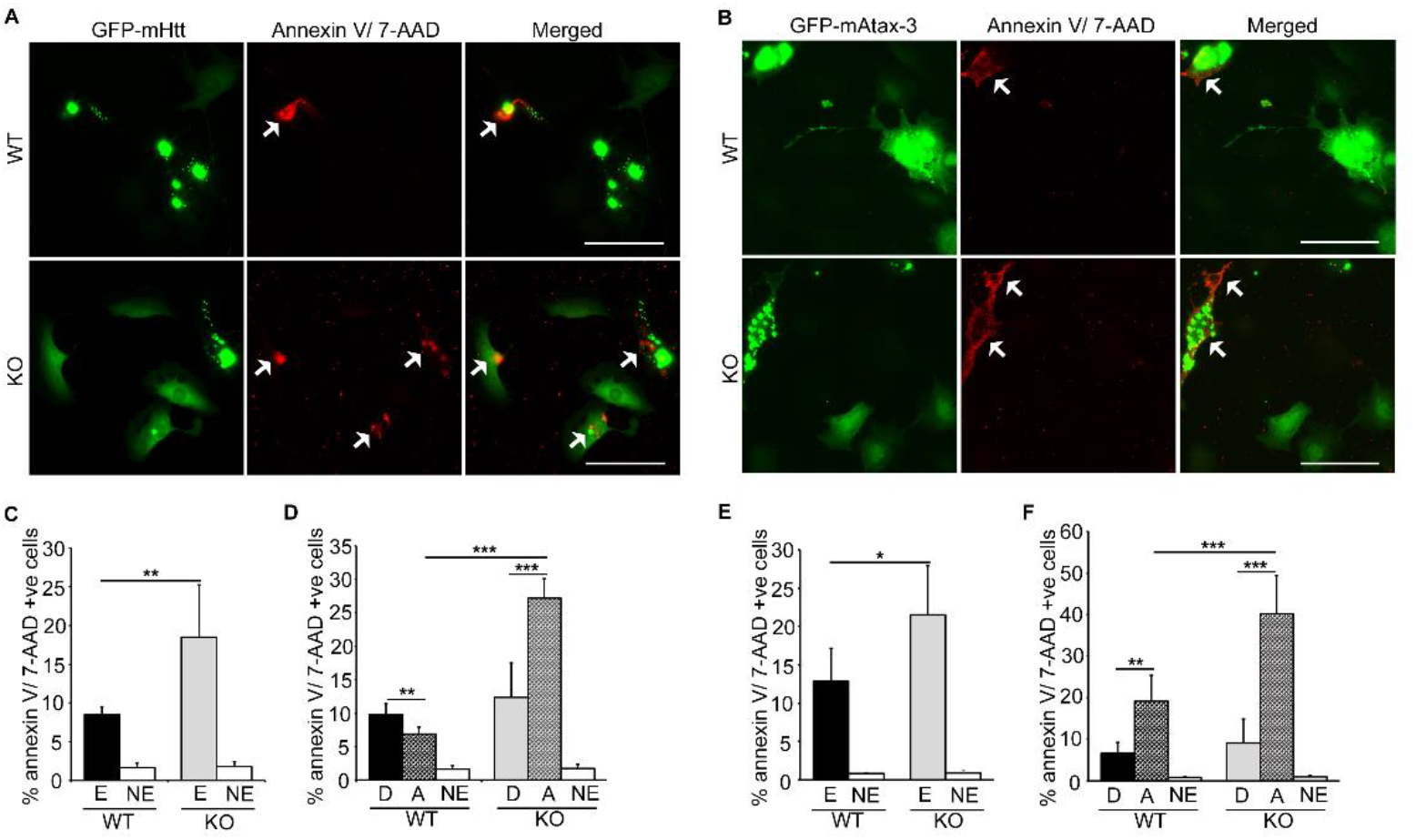
Optineurin deficiency enhances the cytotoxicity caused by mHtt and mAtax-3 aggregates: **(A)** Representative images showing staining for Annexin V/7-AAD in WT and KO (*Optn*^*−/−*^) MEFs expressing GFP-mHtt. Arrows indicate GFP-mHtt expressing cells showing Annexin V/ 7-AAD staining. Scale bar: 100μm. **(B)** Representative images showing staining for Annexin V/ 7-AAD in WT and KO MEFs expressing GFP-mAtax-3. Arrows indicate GFP-mAtax-3 expressing cells showing Annexin V/ 7-AAD staining. Scale bar: 100μm. **(C)** Bar diagram showing the percentage of GFP-mHtt expressing (E) and non-expressing (NE) WT and *Optn*^*−/−*^ MEFs positive for Annexin V/7-AAD. n=6. Bars represent mean ± SD. **p ≤ 0.01. **(D)** Bar diagram showing the percentage of diffused (D) or aggregated (A) GFP-mHtt expressing or non-expressing WT and KO MEFs positive for Annexin V/ 7-AAD. n=6. ***p ≤ 0.001 and **p ≤ 0.01. **(E)** Bar diagram showing the percentage of GFP-mAtax-3 expressing (E) and non-expressing (NE) WT and KO MEFs positive for Annexin V/7-AAD. n=6. *p ≤ 0.05. **(F)** Bar diagram showing the percentage of diffused (D) or aggregated (A) GFP-mAtax-3 expressing and non-expressing WT and KO MEFs positive for Annexin V/ 7-AAD. n=6. ***p ≤ 0.001 and **p ≤ 0.01.

We further analysed the toxic effects caused by expression of mHtt and mAtax-3 in a neuronal cell line, N2A, by knocking down Optn using ShRNA. The ShRNA mediated knock down of Optn was first confirmed by western blotting (Fig. 7A). Upon knockdown of optineurin in N2A cells, the percentage of cells showing mHtt aggregates significantly increased (Fig. 7B and C). This was in accordance with previous observations in Hela (29) and N2A cells (30), which showed that optineurin clears mHtt aggregates by autophagy. Thus, knockdown of optineurin blocked autophagy and the aggregates got accumulated (Fig. 7B and C). We co-transfected N2A cells with GFP-mHtt and optineurin ShRNA (Sh-Optn) or control ShRNA (Sh-Con) and analysed cell death after 48 hours of transfection. We observed that GFP-mHtt caused significantly higher cell death in cells co-transfected with Sh-Optn as compared to those co-transfected with Sh-Con (Fig. 7F and G). Separate death analysis of cells expressing mHtt in non-aggregated and aggregated forms in the control group suggested that non-aggregated form of mHtt was significantly more toxic as compared to the aggregated form (Fig. 7H). When we compared the cell death caused by aggregated mHtt in control and optineurin knock down cells, we observed that Optn knock down cells showed significantly higher cell death (Fig. 7H), suggesting that the aggregates are more toxic in optineurin-deficient cells. Knockdown of optineurin decreased aggregation of mAtax-3 unlike the effect on mHtt aggregates (Fig. 7D and E). This suggests that optineurin promotes sequestration of non-aggregated mAtax-3 protein into aggregates in N2A cells. Optineurin knockdown had no significant effect on total cell death caused by expression of mAtax-3 in N2A cells (Fig. 7J). Consistent with the observation in *Optn*^*−/−*^ MEFs, deficiency of optineurin mediated by knockdown using Sh-Optn in N2A cells increased the toxicity of mAtax-3 aggregates (Fig. 7I and K). These results suggest that optineurin protects the N2A cells from toxicity caused by mutant protein aggregates.

**Figure 7:**
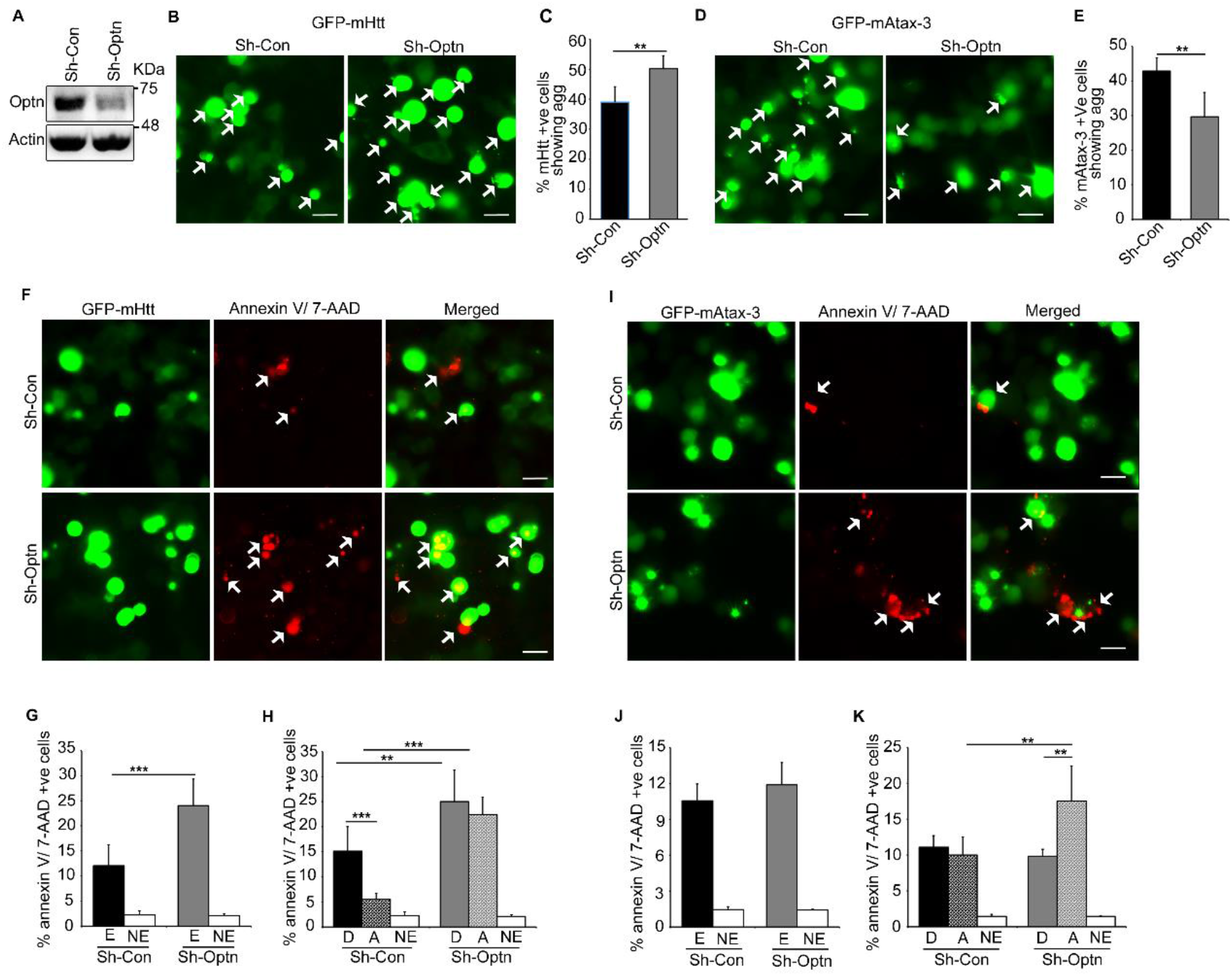
Optineurin knockdown in N2A enhances cell death caused by mHtt and mAtax-3 aggregates: **(A)** Western blot showing knockdown of optineurin mediated by ShRNA in N2A cells after 48 hours of transfection with required plasmids. **(B, D)** Representative images showing GFP-mHtt **(B)** or GFP-mAtax-3 **(D)** expressing cells forming aggregates in N2A cells after 48 hours of co-transfection with control Sh RNA (Sh-Con) or optineurin ShRNA (Sh-Optn). Arrows indicate cells showing aggregates. Scale bar: 20 μm. **(C, E)** Bar diagram showing percentage of GFP-mHtt **(C)** or mHtt **(E)** expressing N2A cells forming aggregates upon co-transfection with Sh-Con or Sh-Optn. n=7 for mHtt and n=6 for mAtax-3. Bars represent mean ± SD. **p ≤ 0.01. **(F)** Representative images of N2A cells showing staining for Annexin V/7-AAD upon co-transfection of GFP-mHtt with Sh-Con or Sh-Optn against optineurin for 48 hours. Arrows indicate mHtt positive cells showing Annexin V/ 7-AAD staining. Scale bar: 20μm. **(G)** Bar diagram showing percentage of GFP-mHtt expressing (E) and non-expressing (NE) N2A cells positive for Annexin V/ 7-AAD upon co-transfection with Sh-Con or Sh-Optn (ShOptn). n=7. ***p ≤ 0.001. **(H)** Bar diagram showing percentage of diffused (D) or aggregated (A) GFP-mHtt expressing and non-expressing N2A cells positive for Annexin V/ 7-AAD upon co-transfection with Sh-Con or Sh-Optn. n=7. ***p ≤ 0.001 and **p ≤ 0.01. **(I)** Representative images of N2A cells showing staining for Annexin V/7-AAD upon co-transfection of GFP-mAtax-3 with Sh-Con or Sh-Optn for 48 hours. Arrows indicate mAtax-3 positive cells showing Annexin V/ 7-AAD staining. Scale bar: 20μm. **(J)** Bar diagram showing percentage of GFP-mAtax-3 expressing and non-expressing N2A cells positive for Annexin V/ 7-AAD upon co-transfection with Sh-Con or Sh-Optn. n=6. **(K)** Bar diagram showing percentage of diffused or aggregated GFP-mAtax-3 expressing and non-expressing N2A cells positive for Annexin V/ 7-AAD upon co-transfection with Sh-Con or Sh-Optn. n=6. **p ≤ 0.01.

## Discussion

We have investigated the role of OPTN in aggregation of two mutant proteins, mHtt and mAtax-3, which are causatively associated with neurodegenerative diseases. Our results suggest that OPTN promotes aggregation of these mutant proteins and this function of OPTN does not require its ubiquitin binding function or LIR, whereas its function in autophagy is dependent on ubiquitin binding and LIR. Therefore, it is clear that this function of OPTN of promoting protein aggregation is independent of its function in autophagy. OPTN promotes aggregation of mAtax-3 in neuronal N2A as well as non-neuronal cells (MEFs). mAtax-3 aggregates do not appear to be cleared by OPTN mediated autophagy in N2A cells or MEFs whereas mHtt aggregates are cleared by OPTN mediated autophagy in neuronal cells but not in MEFs. Thus, it appears that the aggregation promoting function of OPTN is not always linked with subsequent degradation of aggregates by autophagy. Enhanced formation of mutant protein aggregates would lead to a lower level of non-aggregated toxic forms of these mutant proteins in the cell, which is expected to result in reduced toxicity.

The cytotoxicity of mutant proteins that are associated with neurodegeneration, and form aggregates, is primarily due to non-aggregated forms, although aggregates may also be toxic. Our results show that OPTN has a cytoprotective function, which reduces the toxicity caused by aggregates formed by mHtt and mAtax-3 in neuronal and non-neuronal cells. This cytoprotective function of OPTN is independent of its function in autophagy as seen by the protective effect of OPTN on mAtax-3 aggregates, which are not cleared by OPTN mediated autophagy. We suggest that OPTN provides protection from cell death in three different ways-(a) by promoting aggregation of mutant proteins (which would lead to a reduction in the level of more toxic non-aggregated form as seen in case of mHtt), (b) by reducing the toxicity of the misfolded protein aggregates, and (c) by autophagy-dependent clearance of aggregates reported previously. How does Optn provide protection from cytotoxicity of mutant protein aggregates? OPTN may be preventing cell death by forming a shell around aggregates, which may be acting as a physical barrier, thereby altering interaction of aggregated mutant proteins with other cellular proteins. The proposed functions of optineurin in protein aggregation and cytoprotection are shown schematically in Figure 8.

**Figure 8:**
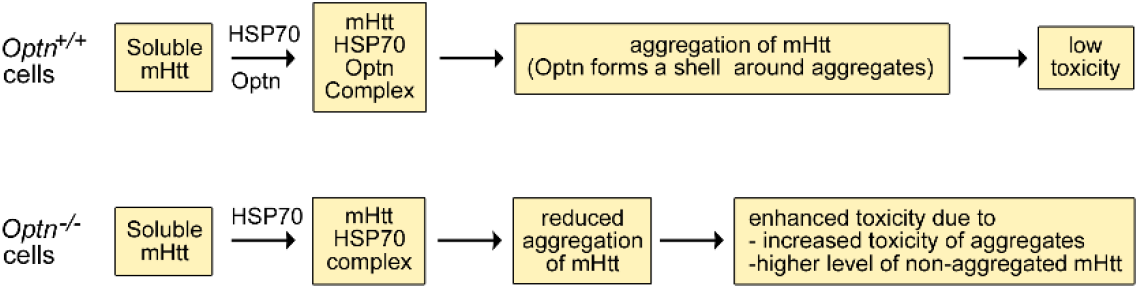
Schematic representation of proposed role of optineurin in promoting mutant protein aggregation and cytoprotection: Mutant protein aggregation into inclusion bodies is a chaperone assisted process. HSP70 (or a related chaperone) is recruited to unfolded/misfolded/ partially folded mHtt (or mAtax-3) protein. In optineurin-positive (wild type) cells, optineurin is also recruited leading to efficient aggregation of mHtt. Optineurin forms a shell around mutant protein aggregates. In optineurin-deficient cells, HSP70 is recruited to unfolded/misfolded/ partially folded mHtt (or mAtax-3) protein, but aggregation of mHtt is less efficient due to lack of optineurin. Enhanced toxicity is seen in optineurin-deficient cells due to two reasons-enhanced toxicity of aggregates and also due to comparatively higher level of non-aggregated mHtt, which is more toxic than aggregated form.

Normal OPTN is observed in pathological structures seen in several neurodegenerative diseases like ALS, Huntington’s disease, Parkinson’s disease, Alzheimer’s disease, Pick’s disease and Creutzfeld-Jacob disease (7, 34). What role optineurin plays in those aggregates is not yet clear. OPTN is involved in autophagy-dependent clearance of some mutant protein aggregates (mutant SOD1 and mHtt), which could be one of the reasons for its association with these aggregates. Our results suggest that the presence of OPTN in these disease-associated aggregates is also likely to be due to its involvement in promoting protein aggregation, and in protecting cells from the toxicity of aggregates.

Mechanism of aggregation of mutant proteins is complex and has been studied extensively. Chaperones of HSP70 family are involved in assisting proper folding of misfolded mutant proteins, and also in their degradation. HSC70/HSP70/HSP90 interact with mutant Htt and influence their aggregation and degradation. Inhibition of HSP90 by chemical inhibitor or knockdown leads to degradation of mHtt and mAtax-3. Heat shock and HSP70 are known to enhance mHtt aggregation and provide cytoprotection (42). Sorbitol, a chemical chaperone has been shown to promote mHtt aggregation and reduce cytotoxicity of mHtt. We observed that the C-terminal domain of OPTN, which is sufficient for promoting mutant protein aggregation, is able to interact with HSC70/HSP70/HSP90. We also observed that inhibition of Hsp90 by a chemical inhibitor, which induces HSP70, enhanced mutant protein aggregation. These observations indicate that OPTN might be modulating the function of HSP70 (or a related chaperone) to promote mutant protein aggregation.

Overall, our results show that OPTN promotes formation of mutant protein aggregates and this function of OPTN is expected to aid in cell survival due to a reduction in toxic non-aggregated forms. Our work also shows that OPTN reduces cytotoxicity caused by mutant protein aggregates, possibly by forming a shell around them. Presence of OPTN in the pathological structures containing protein aggregates seen in several neurodegenerative diseases could be due to its involvement in protein aggregation, and in protecting cells from the toxic effects of protein aggregates.

## Materials and Methods

### Plasmid constructs

HA-OPTN, HA-E478G-OPTN and the deletion constructs HA-1-412-OPTN and HA-412-577-OPTN have been described earlier (46). Mouse Optn ShRNA was described in (47). GFP-mHtt 97Q was described in (36). GFP-mAtax-3 130Q was kindly provided by Dr. Nihar Ranjan Jana (43).

### Antibodies and Reagents

Following primary antibodies were used: mouse monoclonal anti GFP (Santacruz; Cat. No. sc-9996; WB 1:1000), mouse monoclonal anti OPTN (Santacruz; Cat. No. sc-166576; WB 1:500), mouse monoclonal anti actin (Millipore; Cat. No. MAB1501; WB 1:10,000), rabbit monoclonal anti HA (CST; Cat. No. 3724; WB 1:1000; IF 1:800), mouse monoclonal anti HA (CST; Cat. No. 2367; WB 1:1000; IF 1:1000), mouse monoclonal anti ubiquitin (Santacruz; sc-8017; IF 1:100), rabbit polyclonal anti p62 (Sigma; WB: 1:1000; IF:1:200). The reagents used were chloroquine (Sigma; Cat. No. C6628), 17-AAG (Santacruz; sc-200641).

### Cell culture and transfections

*Optn*^*−/−*^ and wild type MEFs have been described (36, 44). MEFs and N2A cells were maintained in Dulbecco’s Modified Eagle Medium (DMEM) supplemented with 10% FBS, 50 IU/ml penicillin and 50 μg/ml streptomycin in a humidified 5% CO2 incubator at 37°C. Cells were plated at appropriate density and transfections with desired plasmids were performed with Jet PRIME (Polyplus transfection; Cat. No. 114) or Lipofectamine 2000 (Invitrogen; Cat. No. 11668027) transfection reagents according to manufacturer’s protocols.

### Immunofluorescence staining and microscopy

Cells were seeded at desired cell density on glass coverslips and after 24 hours, transfections with required plasmids were performed. The cells were fixed with 4% formaldehyde after 24-48 hours of transfections and immunostaining with desired antibodies was carried out as described (36, 44). Coverslips with stained cells were then mounted on glass slides in mounting medium containing DAPI. They were then visualized under a fluorescent microscope for counting percent aggregation or cell death. To scan confocal images, Leica TCS SP8 confocal microscope with 63 X oil immersion objective and 1.4 NA was used. Optical sections of 0.4 μm thickness were taken.

### Aggregation assay

WT and *Optn*^*−/−*^ MEFs or N2A cells were seeded on coverslips. Next day, transfections with either GFP-mHtt 97Q or GFP-mAtax-3 130Q plasmids were performed. 48 hours post transfection, cells were fixed and analysed under a fluorescent microscope for percent of GFP-mHtt 97Q or GFP-mAtax-3 130Q positive cells forming aggregates (29, 30). At least 200 expressing cells were counted in each experiment and each experiment was repeated the indicated number of times. In the experiments in which non-aggregated and aggregated GFP expressing cells were analysed separately, at least 200 cells in each group were counted.

### Cell death assay

For cell death analysis, GFP Certified Annexin V/ 7-AAD staining kit (Enzo, ENZ-51002-100) was used as described earlier (36). Briefly, 48 hours post transfection, cells were washed thrice with PBS and stained with Annexin V and 7-AAD (7 aminoactinomycin D) for 15 minutes in dark at room temperature. They were then washed and fixed with 2% formaldehyde and mounted in media containing DAPI. These cells were then observed under a fluorescent microscope for dead cells staining positive for Annexin V binding to the exposed phosphatidyl serine and 7-AAD binding to fragmented DNA. Percentage of total GFP-mHtt 97Q or GFP-mAtax-3 130Q positive cells staining for annexin V/ 7-AAD was analysed. Segregated cell death analysis was also done for cells having non-aggregated or aggregated GFP-mHtt 97Q or GFP-mAtax-3 130Q. At least 200 expressing cells were counted in each experiment and experiments were repeated as indicated in figure legends. In the experiments in which non-aggregated and aggregated GFP expressing cells were analysed separately, at least 200 cells in each group were counted.

### Fractionation and insolubility assay

Cells were lysed in NP-40 lysis buffer (50 mM Tris, 150 mM NaCl, 1 mM EDTA, 5mM MgCl2, 0.5% NP-40, pH 7.4, supplemented with Complete Protease Inhibitor Cocktail, Roche). After centrifugation at 5000g for 10minutes, the supernatant was aspirated and used as soluble fraction. The pellet was washed twice and re-suspended in NP-40 buffer and was used as insoluble fraction. Both soluble and insoluble fractions were then boiled in SDS sample buffer for 10 minutes and loaded on SDS polyacrylamide gel and western blotting was performed (29, 30). ImageJ software was used to quantitate the band intensity by densitometry method. Actin was taken as loading control. Insolubility index was calculated by taking the ratio of intensity of band in insoluble fraction to that in soluble fraction.

### Statistical analysis

Quantitation of data is presented mean ± SD (standard deviation). Error bars represent SD. Significance of difference is calculated by using Student’s two tailed T test.

## Acknowledgements

GS acknowledges the J.C. Bose National Fellowship grant (SR/S2/JCB-41/2010) from the Department of Science and Technology, Government of India. SCM acknowledges the Council for Scientific and Industrial Research, India for research fellowship.

## Author contributions

SCM and AKR performed the experiments. GS and SCM planned the experiments, analyzed the data and wrote the paper. GS conceived and obtained funding for this study. All authors approved final version of the manuscript

## Notes

### Competing Interest Statement

The authors have declared no competing interest.

