## Supplementary Figures 1,2 for "Optineurin promotes aggregation of mutant huntingtin and mutant ataxin-3, and reduces cytotoxicity of aggregates"

***Correspondence to:**

Ghanshyam Swarup

**This file includes:**

Figures S1 and S2

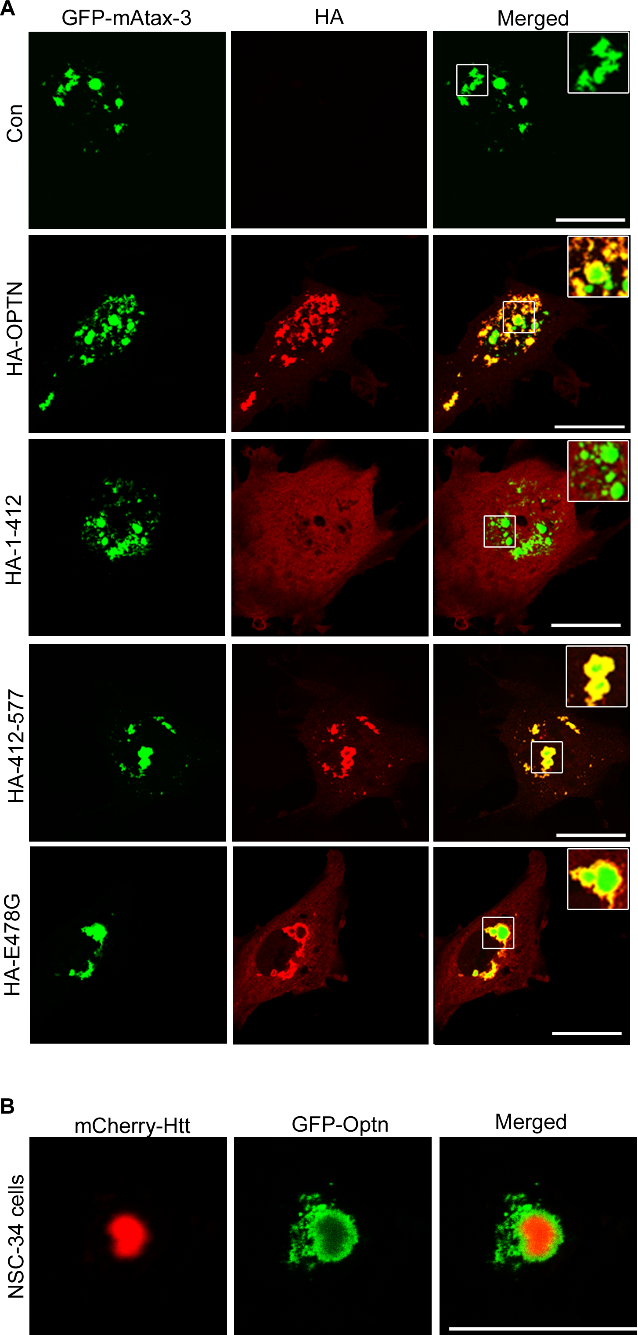

Fig. S1: Localization of optineurin around mutant protein aggregates: (A) Confocal microscopy images showing HA-OPTN, HA-412-577-OPTN and E478G-OPTN form a shell around GFP-mAtax-3 aggregates in *Optn^-/-^* MEFs whereas HA-1-412-OPTN does not. Scale bar: 25μm. Three confocal sections are merged in the images. (B) Confocal microscopy image showing GFP-Optn forms a shell around mCherry-mHtt aggregate in NSC-34 cells. Scale bar: 25μm. Three confocal sections are merged in the images.

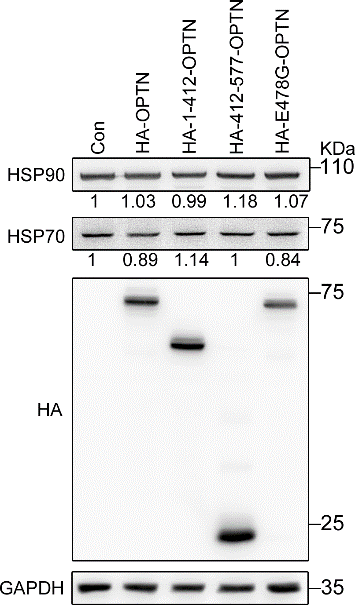

**Fig. S2**: Western blot showing the levels of HSP90 and HSP70 upon expression of different mutants of optineurin in *Optn^-/-^* MEFs. ‘Con’ represents control pCDNA transfected samples.
